# A partner switch from CDS2 to RAB1A redirects MBOAT7 to inhibit DGAT2-mediated lipid droplet growth

**DOI:** 10.1101/2025.08.26.672501

**Authors:** Jiesi Xu, Jianxin Zhang, Wei Wang, Honggang Su, Siyu Chen, Mei Ding, Xun Huang

## Abstract

Metabolic dysfunction-associated steatotic liver disease (MASLD) is a highly prevalent disorder worldwide characterized by the accumulation of hepatic lipid droplets (LDs). However, the mechanisms by which hepatocytes adapt to the dynamic changes in LD morphology to maintain metabolic homeostasis remains incompletely understood. Here, we show that MBOAT7, a known MASLD-associated protein, cooperates with CDS2 to regulate lipid droplet dynamics and lipid metabolism. Under physiological conditions, their interaction in the endoplasmic reticulum (ER) maintains metabolic homeostasis. Disruption of this interaction, such as through CDS2 knockdown or loss of function, triggers an adaptive response wherein MBOAT7 translocates to ER-LD contact sites in a RAB1-dependent manner. This redistribution inhibits DGAT2-mediated LD growth and promotes lipolysis. Our findings highlight the redistribution of MBOAT7 to ER-LD contacts serves as a critical mechanism for adaptive control LD size and lipid metabolic homeostasis.

## Introduction

The liver serves as the central hub for systemic lipid metabolism, orchestrating the complex processes of fatty acid synthesis, storage, and mobilization, as well as cholesterol homeostasis. Disruptions in this delicate equilibrium are a hallmark of metabolic dysfunction-associated steatotic liver disease (MASLD), a burgeoning global health concern that can progress from simple hepatic steatosis to steatohepatitis (MASH), fibrosis, and cirrhosis (1).

The excessive accumulation of hepatic lipid droplets (LDs) is a central pathological feature of MASLD (2). As dynamic organelles, LDs serve as key hubs for neutral lipid storage and metabolism, and their formation acts as a compensatory mechanism to protect hepatocytes from cytotoxic free fatty acids and other bioactive lipids. However, while initially adaptive, persistent LD expansion causes physical distortion of hepatocytes, nuclear displacement, and alterations in liver parenchymal architecture (3–5). Although the triggers and consequences of hepatic LD accumulation in MASLD have been extensively studied, how hepatocytes actively counteract and mitigate the increasing LD expansion remains incompletely understood. Two major adaptive strategies have emerged: first, lipolysis and lipophagy, which preferentially target smaller LDs that are more accessible for degradation (6, 7); and second, remodeling of inter-organelle contacts between LDs and other organelles—such as the endoplasmic reticulum, mitochondria, and lysosomes, facilitating lipid processing and degradation (8, 9). Through these mechanisms, LD remodeling serves as a vital process for maintaining metabolic homeostasis. LD-associated proteins are known to regulator of LD size, yet their specific roles in counteracting LD expansion await further elucidation.

A critical, yet often overlooked, aspect of hepatic lipid regulation is the spatial organization of metabolic enzymes within cellular organelles and the dynamic contact sites between them, particularly those between the endoplasmic reticulum (ER) and lipid droplets (LDs). These ER-LD contact sites are emerging as vital platforms for lipid trafficking, synthesis, and signaling (8, 10–13), but the molecular mechanisms that govern protein targeting to these sites and their functional consequences for whole-body metabolism remain incompletely understood.

Membrane Bound O-Acyltransferase Domain Containing 7 (MBOAT7) and CDP-Diacylglycerol Synthase (CDS) are two enzymes implicated in distinct branches of phospholipid metabolism. MBOAT7, an ER-resident enzyme, incorporates polyunsaturated fatty acids into lysophosphatidylinositol through the PI remodeling Lands’ cycle (14), and its loss-of-function variants in humans are strongly associated with an increased risk of MASLD, liver fibrosis, and hepatocellular carcinoma (15–18). Parallel to this pathway, PI can also be synthesized *de novo* via the Kennedy pathway, in which the conversion of phosphatidic acid to CDP-diacylglycerol (CDP-DAG) by CDS (CDP-DAG synthase) constitutes a critical step (19, 20). Notably, CDS2, one of the two CDS proteins in mammals, is selectively inhibited by arachidonoyl-containing PI species (PI-AA), suggesting a regulatory feedback loop that links MBOAT7-mediated PI remodeling with CDS2-mediated *de novo* PI synthesis (21). This interplay is biologically significant: *Mboat7* deficiency has been shown to promote hepatic lipid accumulation and steatohepatitis, partly through increasing CDS2 activity (22, 23). Conversely, deletion of *Cds2* leads to the formation of large LDs and spontaneous development of metabolic dysfunction-associated steatohepatitis (MASH) (24–26), providing a valuable model for studying hepatocyte adaptations to drastic LD morphological changes. While the individual roles of these enzymes in their respective biochemical pathways are recognized, a broader, integrative understanding of how they might functionally interact to coordinate hepatic lipid flux has been lacking.

Here, we uncover a coordinated mechanism by which MBOAT7 and CDS2 jointly regulate LD dynamics. We demonstrate that *CDS2* deletion triggers the adaptive relocation of MBOAT7 to ER-LD contact sites, where it restricts LD expansion by competing with RAB1-mediated DGAT2 targeting to LD. These findings establish MBOAT7 and CDS2 as key regulators of a membrane-metabolic crosstalk that integrates inter-organelle communication with lipid homeostasis, revealing a robust adaptive response to hepatic steatosis in hepatocytes.

## Results

### MBOAT7 is localized to the ER-LD contacts in lipid-loaded *CDS2* KO cells

We have previously shown that *Cds2* deficiency causes a rapid development of MASH (26). To explore the cellular function of CDS2, we conducted APEX2-based proximity labeling using an APEX2-CDS2 fusion protein to identify its proximal partners (Figure S1A). Across two biological replicate experiments, we identified 24 proteins specifically enriched in APEX2-CDS2-expressing cells compared to APEX2 alone control. Given the ER localization of CDS2, we focused on ER-associated proteins (Figure S1B). Among these, MBOAT7 emerged as a protein of particular interest due to its established role in phosphatidylinositol biosynthesis and its associated with MASLD. Therefore, we chose MBOAT7 for further investigation.

To investigate the functional interplay between MBOAT7 and CDS2, we first studied whether their enzymatic activities were affected by each other. Previous report show that the activity of CDS2 is selectively inhibited by arachidonoyl-containing PI species (PI-AA), the product of MBOAT7-mediated PI remodeling (21). Therefore, we measured whether CDS2 affects the acyltransferase activity of MBOAT7 using MBOAT7 substrates, lyso-PI and [14C]-arachidonic acid. We analyzed the activity of MBOAT7 in the ER from the liver of control mice (*Cds2^f/f^*), mice with liver-specific depletion of *Cds2* (*Cds2^f/f^;AlbCre*), and hepatic *Cds2*-depleted mice with AAV-mediated hepatic re-expression of *Cds2* (*Cds2^f/f^;AlbCre* + AAV-TBG-*Cds2*). The results showed that *Cds2* deficiency did not significantly impact the enzymatic activity of MBOAT7 (Figure 1A).

**Figure 1.**
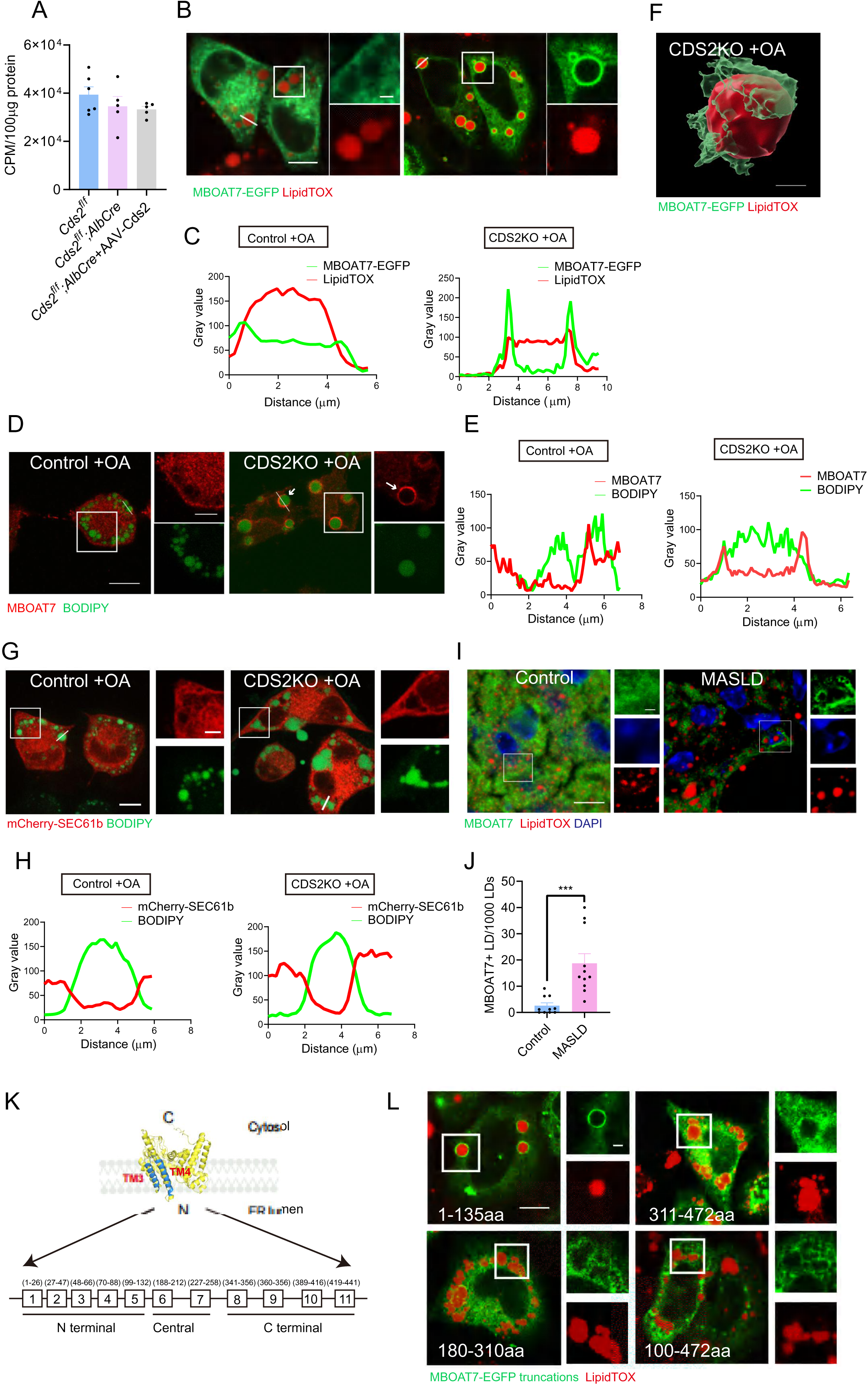
MBOAT7 is localized to the ER-LD contacts in CDS2 KO cells. (A) MBOAT7 enzymatic activity was measured in *Cds2^f/f^*, *Cds2^f/f^;AlbCre* mice, and *Cds2^f/f^;AlbCre* mice injected with AAV-TBG-*Cds2*. N=5-6 mice per group. (B) *CDS2* KO and control HeLa cells were transfected with MBOAT7-EGFP and treated with 100 μM OA for 12 hours, followed by LD labeling with LipidTOX. (C) Fluorescence colocalization was analyzed along the white line in the main confocal image. (D) *CDS2* KO and control HeLa cells were treated with 100 μM OA for 12 hours and immunostained with anti-MBOAT7 antibody. White arrows indicate nicks on LD surface. (E) Fluorescence colocalization was analyzed along the white line in the main confocal image. (F) A 3D-SIM image reveals that MBOAT7-EGFP enwraps a LD in an OA-treated *CDS2* KO cell. Scale bar: 2 μm. (G) *CDS2* KO and control HeLa cells were transfected with mCherry-SEC61b (an ER marker), then treated with 100 μM OA for 12 hours. LDs were labeled with BODIPY. (H) Fluorescence colocalization was analyzed along the white line in the main confocal image. (I) C56BL/6 mice were fed a chow (control) or GAN (MASLD) diet (40 kcal% fat, 20 kcal% fructose and 2% cholesterol) for 26 weeks. Liver sections were immunostained with anti-MBOAT7 antibody. White arrows indicate MBOAT7 enclosing LDs. (J) The number of LDs surrounded by MBOAT7 was quantified by analyzing 1,000 LDs in confocal images. N=9-11 liver sections per group. (K) Top: 3D structure of MBOAT7 in the ER membrane, highlighting its transmembrane (TM) domains. Bottom: The 11 TM domains of MBOAT7 (boxes) and the three regions of the protein expressed as truncations. (L) *CDS2* KO cells transfected with truncated versions of MBOAT7 were treated with 100 μM OA for 12 hours, followed by LD labeling with LipidTOX. Except for (F), scale bars represent 10 μm in main images and 2 μm in enlarged images.

Previous studies have shown that *CDS2* knockout induced large LDs (25). Given the shared subcellular localization and functional interplay between MBOAT7 and CDS2, we sought to investigate the localization and roles of MBOAT7 in LD regulation. MBOAT7-EGFP exhibited an ER-like localization pattern in both wild-type and *CDS2* KO HeLa cells (Figure S2A and B). Interestingly, in *CDS2* KO cells treated with oleic acid (OA) to induce LD formation, MBOAT7-EGFP surrounded LDs stained by LipidTOX (Figure 1B-C). In contrast, MBOAT7-EGFP did not show colocalization with LDs in OA-treated control HeLa cells (Figure 1B-C). Furthermore, in OA-treated *CDS2* KO cells, MBOAT7-EGFP was colocalized with the LD marker mScarlet-PLIN1, whereas no such colocalization was observed in control cells (Figure S2C). Similarly, in OA-treated HepG2 cells, MBOAT7-EGFP remained localized to the ER, but enclosed LDs when *CDS2* was knocked down (Figure S1D). To rule out the possibility that the phenomena is caused by EGFP tagging, we also analyzed the localization of endogenous MBOAT7. In OA-treated *CDS2* KO cells, endogenous MBOAT7 colocalized with LDs, but this was not observed in OA-treated control cells (Figure 1D-E). We also notice that under these conditions, the MBOAT7 protein surrounding the LD surface is often incomplete, with one or more nicks (Figure 1D arrow).

There are several mechanisms for protein localization on the LDs, including short hairpin, hydrophobic region (which may form an amphipathic helix), and lipid modification (27). Since MBOAT7 is a multipass transmembrane protein, it is likely localized to ER regions adjacent to LDs rather than directly to the LD phospholipid monolayer. Indeed, 3D reconstruction of the LDs confirmed that MBOAT7 partially surrounds the LDs in OA-treated *CDS2* KO cells (Figure 1F).

The observed targeting of ER-localized MBOAT7 to the ER-LD contacts in *CDS2* KO cells prompted us to investigate whether *CDS2* depletion affects the targeting of other ER proteins to LDs. The ER marker mScarlet-SEC61b did not colocalize with LDs in OA-loaded *CDS2* KO HeLa cells (Figure 1G-H). Next, we examine whether this relocalization of MBOAT7 to ER-LD is related to MASH pathogenesis. Since CDS2 protein levels are downregulated in MASH mouse models (26), we also examined MBOAT7 localization in the liver of the fat/fructose/cholesterol-rich Gubra-Amylin NASH (GAN) diet feeding-induced MASH mice. Confocal imaging showed that endogenous MBOAT7 was more enriched at the LD periphery in liver sections from GAN diet-induced MASH mice compared to those from normal mice (Figure 1I-J). These findings suggest that MBOAT7 specifically localizes to the ER-LD contacts in lipid-overloaded *CDS2*-deficient cells but not in control cells.

MBOAT7 consists of 11 transmembrane (TM) helices organized into two bundles (TM1-5 and TM9-11) that tilt apart, creating a central catalytic chamber (28) (Figure 1K). To identify the region required for the localization of MBOAT7 to the ER-LD contacts, we generated three EGFP-tagged truncated versions of MBOAT7, containing the N-terminal region (amino acids 1-135), the central region (180–310), and the C-terminal region (311–472) (Figure 1K). Only the N-terminal region localized to the ER-LD contacts in *CDS2* KO cells, and MBOAT7 with a deletion of the N-terminal 99 amino acids did not localize to the ER-LD contacts (Figure 1L).

Therefore, the N-terminus of MBOAT7 is required for the localization of MBOAT7 to ER-LD contacts in *CDS2* KO cells.

### MBOAT7 interacts with CDS2

Next, we investigated the mechanism underlying MBOAT7 localization to ER-LD contact sites specifically in *CDS2* KO cells. Since both CDS2 and MBOAT7 are localized to the ER and MAMs (26) and MBOAT7 is found in CDS2 proximal proteome, we hypothesized that CDS2 retains MBOAT7 within the ER, while MBOAT7 relocates to the ER-LD contacts upon *CDS2* depletion. To test this, we performed co-immunoprecipitation (co-IP) assays, which show that Flag-CDS2 precipitated MBOAT7-EGFP (Figure 2A), and, reciprocally, MBOAT7-EGFP precipitated Flag-CDS2 (Figure 2B). To rule out potential artifacts from protein overexpression, we used CRISPR-based knock-in (KI) to tag the endogenous CDS2 locus with an N-terminal EGFP and confirmed that endogenous CDS2 coprecipitated with endogenous MBOAT7 in HeLa cells (Figure 2C). To examine the direct binding between these two proteins, we purified His-MBOAT7 from E. coli and Flag-CDS2 from mammalian cells. Western blot analysis confirmed their direct interaction (Figure 2D). Collectively, these results demonstrate the interaction between CDS2 and MBOAT7, which retains MBOAT7 at the ER.

**Figure 2.**
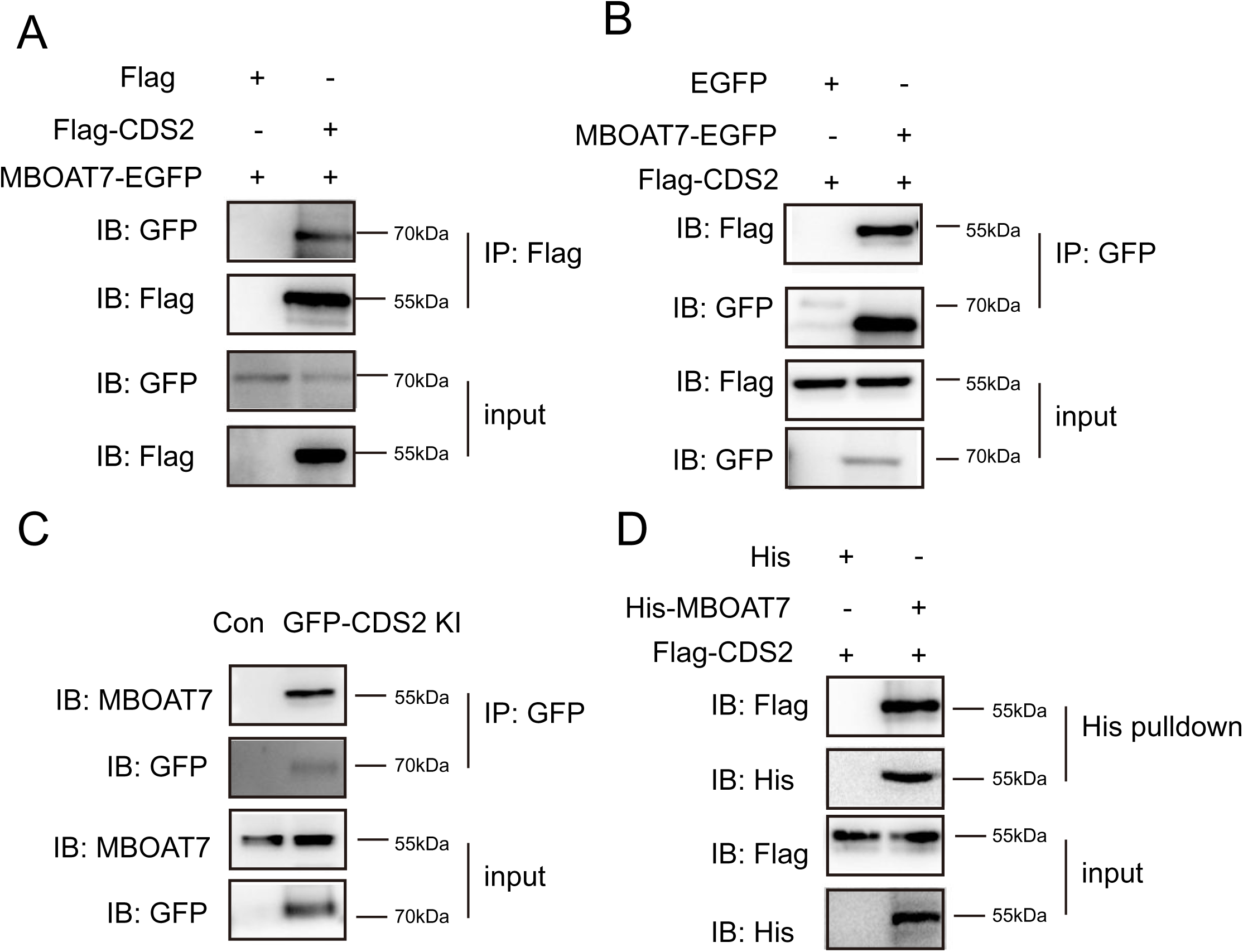
MBOAT7 interacts with CDS2. (A) Co-IP assay showing that Flag-CDS2 precipitates MBOAT7-EGFP. (B) Co-IP assay showing that MBOAT7-EGFP precipitates Flag-CDS2. (C) Co-IP assay demonstrating that endogenous CDS2 precipitates MBOAT7. (D) *In vitro* pull-down assay indicating that Flag-CDS2 directly binds to MBOAT7-His.

### RAB1A regulates the localization of MBOAT7 at the ER-LD contacts in *CDS2* KO cells

Next, we investigated how MBOAT7 is recruited to the ER-LD contacts in *CDS2* KO cells. ER proteins typically target the LD monolayer surface, which requires membrane continuity between the ER outer leaflet and the LD monolayer (11). This targeting can occur either during early LD formation or to mature LDs after their formation. These ER proteins associate with the LD monolayer through hydrophobic hairpins, which insert midway into the lipid bilayers of the ER, with both the N- and C-termini facing the cytosol (29). Since MBOAT7 lacks a hairpin structure, we hypothesized that ER-localized MBOAT7 associates with LDs in *CDS2* KO cells via interaction with LD-associated proteins.

We conducted an unbiased proteomic screen for MBOAT7-interacting proteins in *CDS2* KO Hela cells. We immunoprecipitated MBOAT7-EGFP from control and *CDS2* KO cells and identified its interacting proteins using LC-MS/MS (Figure 3A). We hypothesized that MBOAT7 would interact with LD proteins in *CDS2* KO cells but not in control cells, so we selected candidates whose interaction with MBOAT7 in *CDS2* KO cells was at least 2-fold stronger than in control cells. Based on previously published LD proteomes (30–32), we focused on LD proteins. Five proteins from our dataset were shared with all the published LD proteomes (Figure 3B). Interestingly, of these five candidates, four of them are RAB proteins (RAB1A, RAB7A, RAB14 and RAB18) (Figure 3C). Previous studies reported that RAB1 mediates targeting triglyceride biosynthetic enzymes to LDs (13, 33). Our results showed that knockdown of *RAB1A* impaired the LD targeting of MBOAT7 in OA-loaded *CDS2* KO cells (Figure 3D-F). To determine whether this regulatory relationship involves a direct physical interaction, co-IP assays showed that MBOAT7-EGFP interacted with Flag-RAB1A in *CDS2* KO cells but not in control cells, suggesting that RAB1A helps MBOAT7 target to the ER-LD contacts in *CDS2* KO cells (Figure 3G). Furthermore, expressing HA-tagged CDS2 in *CDS2* KO cells inhibited the interaction between MBOAT7 and RAB1A (Figure 3H).

**Figure 3.**
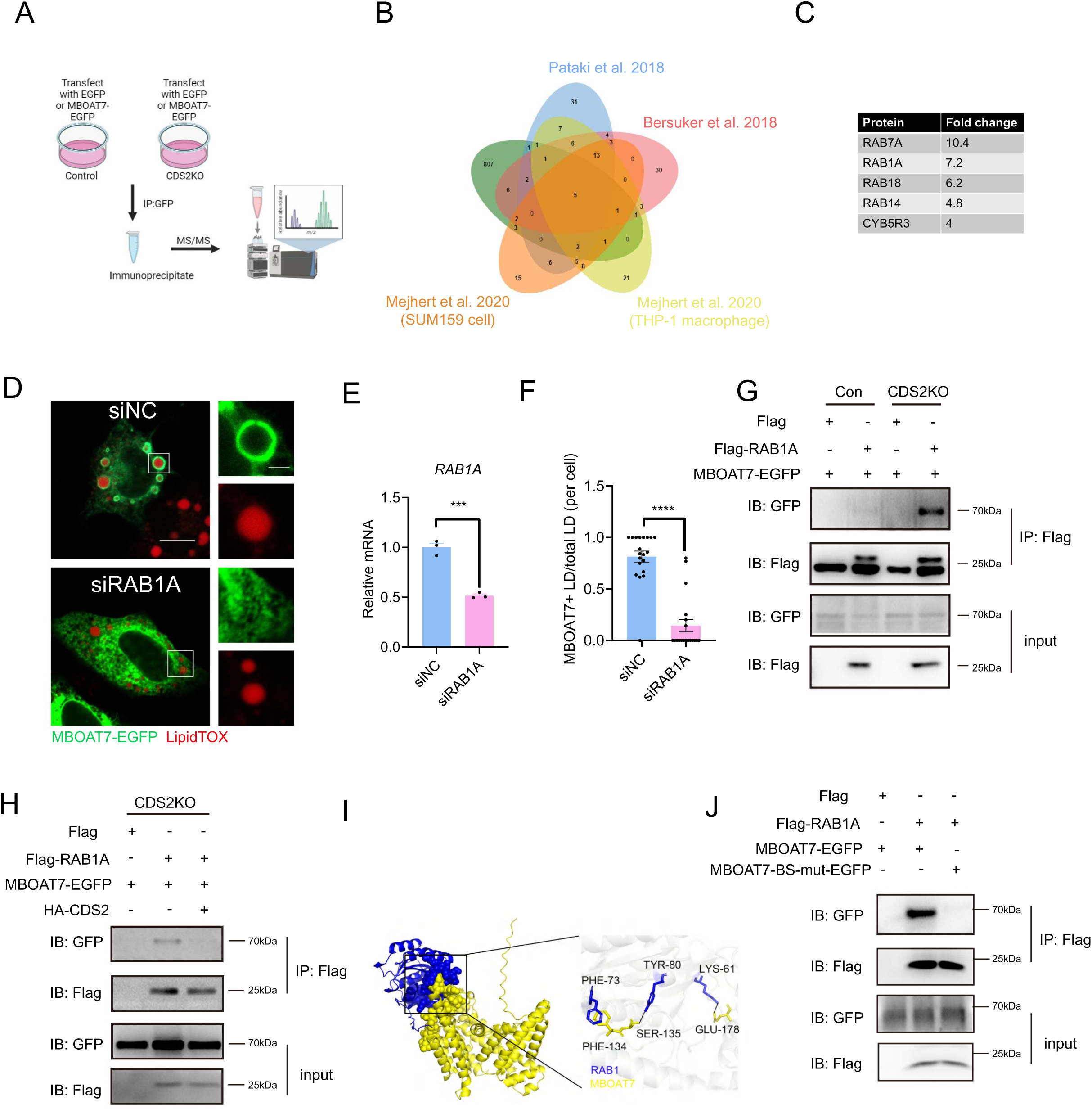
RAB1A is required for targeting of MBOAT7 to LDs in *CDS2* KO cells. (A) Schematic of the workflow for identifying MBOAT7-interacting proteins in control and *CDS2* KO cells. (B) Venn diagram illustrating the overlap between the MBOAT7-interacting proteins from (A) and LD proteins from various LD proteomes. (C) Five MBOAT7-interacting proteins identified from the Venn diagram in (B). (D) *CDS2* KO cells were transfected with siNC or siRAB1A, and MBOAT7-EGFP, then treated with 100 μM OA for 12 hours. LDs were labeled with LipidTOX. (E) *RAB1A* mRNA levels in HeLa cells transfected with control siRNA or siRAB1A. (F) Quantification of LDs surrounded by MBOAT7-EGFP in control (siNC) or RAB1A knockdown (siRAB1A) cells. (G) Co-IP assay showing that Flag-RAB1A precipitates MBOAT7-EGFP in *CDS2* KO cells. (H) Co-IP assay showing that Flag-RAB1A precipitates MBOAT7-EGFP in *CDS2* KO cells, and expression of HA-CDS2 in *CDS2* KO cells disrupts the RAB1A/MBOAT7 interaction. (I) 3D structure of MBOAT7/RAB1A docking and the potential binding sites. (J) Co-IP assay showing that Flag-RAB1A precipitates MBOAT7-EGFP, but fails to interact with MBOAT7-BS-mut-EGFP, in which the RAB1A-binding sites are mutated in *CDS2* KO cells. *** p < 0.001, **** p < 0.0001.

We then determined the binding sites between MBOAT7 and RAB1A. MBOAT7 structure were shown previously (28). The crystal structure of RAB1A was predicted using AlphaFold3, and the 3D structure of MBOAT7/RAB1A docking was simulated. The results showed that the amino acid residues F134, S135, E178 in MBOAT7 and the amino acid residues F73, Y80, K90 in RAB1A were the potential interaction sites (Figure 3I). Mutation of the predicted binding sites within MBOAT7 to alanine abrogated their interaction (Figure 3J). Together, these results support a model of a regulated partner switch: CDS2 retains MBOAT7 in the ER by competing with RAB1A, but upon *CDS2* depletion, RAB1A binds MBOAT7 and recruits it to ER-LD contact sites.

### MBOAT7 reduces LD size in *CDS2* KO cells by inhibiting the targeting of DGAT2 to LDs

We then investigated the cellular and physiological functions of ER-LD contact-localized MBOAT7. Overexpression of MBOAT7-EGFP did not affect LD size in control cells, but it reduced LD size in *CDS2* KO cells (Figure 4A-B). Overexpression of an MBOAT7 mutant with reduced catalytic activity (MBOAT7(N321A, H356A)-EGFP) still reduced the LD size in *CDS2* KO cells (Figure 4A-B). This indicates that the effect of MBOAT7 on LD size reduction in *CDS2* KO cells is independent of its enzymatic activity.

**Figure 4.**
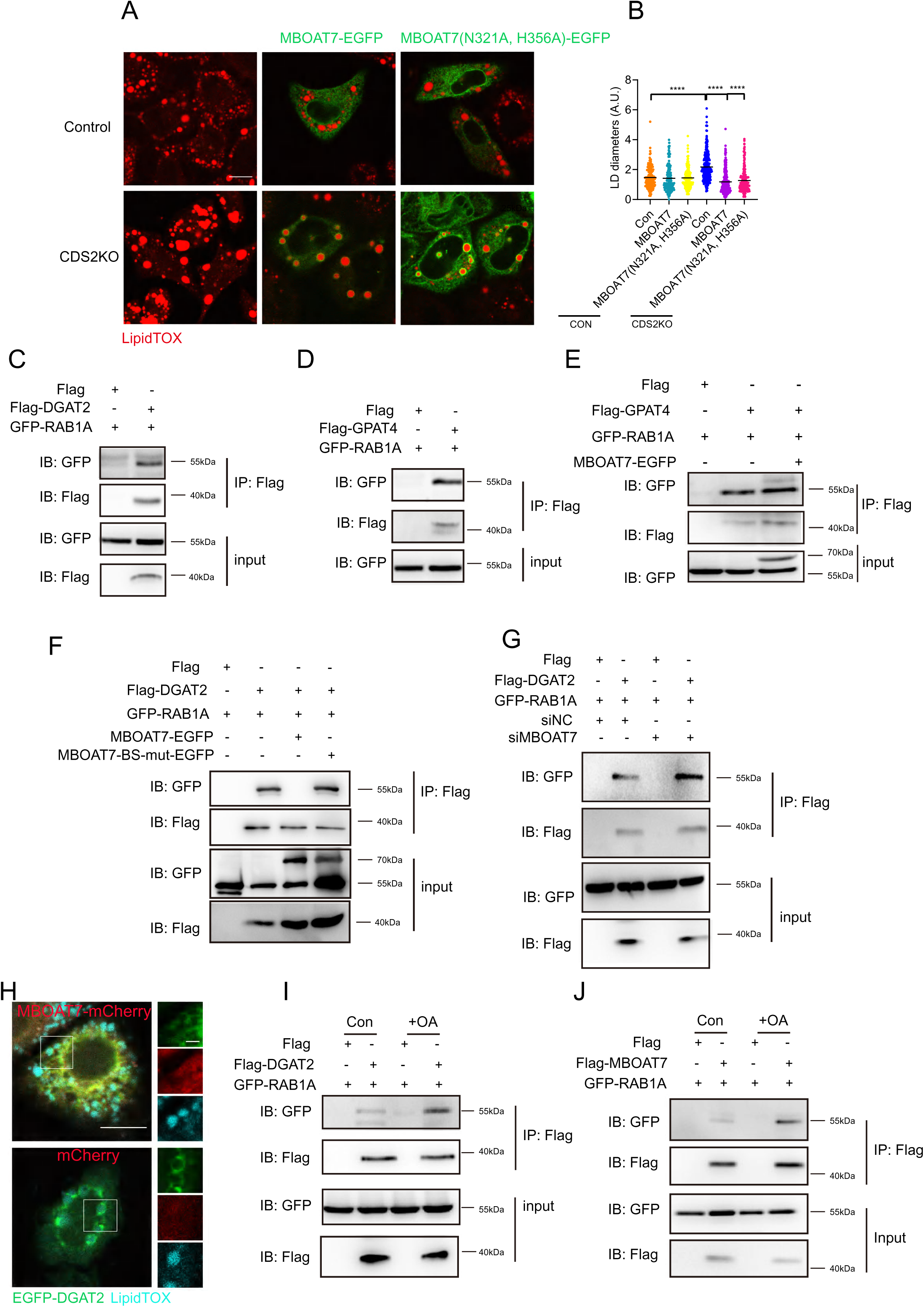
MBOAT7 reduces LD size in *CDS2* KO cells by competing with DGAT2 for targeting to LDs. (A) Control HeLa or *CDS2* KO cells were transfected with MBOAT7-EGFP or the MBOAT7(N321A, H356A)-EGFP mutant, then treated with 100 μM OA for 12 hours. LDs were labeled by LipidTOX. Scale bar: 10 μm. (B) Quantification of LD sizes from (A). N=200 per group. (C) 293T cells were transfected with GFP-RAB1A and Flag-DGAT2 or Flag control plasmids. Co-IP assay shows that Flag-DGAT2 precipitates GFP-RAB1A. (D) 293T cells were transfected with GFP-RAB1A and Flag-GPAT4 or Flag control plasmid. Co-IP assay shows that Flag-GPAT4 precipitates GFP-RAB1A. (E) *CDS2* KO cells were transfected with GFP-RAB1A and Flag-GPAT4 with or without MBOAT7-EGFP. Co-IP shows that Flag-GPAT4 precipitates GFP-RAB1A. (F) *CDS2* KO cells were transfected with GFP-RAB1A and Flag-DAGT2 with or without MBOAT7-EGFP or MBOAT7-BS-mut-EGFP. Co-IP shows that Flag-DGAT2 precipitates GFP-RAB1A. (G) *CDS2* KO cells were transfected with Flag-DGAT2 and GFP-RAB1A with siRNA control (siCon) or siMBOAT7. Co-IP assay shows that Flag-DGAT2 precipitates GFP-RAB1A. (H) Confocal images showing that EGFP-DGAT2 targets to LDs in *CDS2* KO cells. MBOAT7-mCherry expression blocks LD targeting of DGAT2. Scale bar: 10 μm in main images and 2 μm in enlarged images. (I) Co-IP assay showing that Flag-DGAT2 precipitates GFP-RAB1A in *CDS2* KO cells treated with or without OA. (J) Co-IP assay showing that Flag-MBOAT7 precipitates GFP-RAB1A in *CDS2* KO cells treated with or without OA. **** p < 0.0001.

To explore the underlying mechanism, we considered previous findings suggesting that the lipid biosynthetic enzymes DGAT2 and GPAT4 target LDs earlier in *CDS2* KO cells than in control cells, promoting LD growth (24). Knockdown of DGAT2 or GPAT4 reduced the LD size in *CDS2* KO cells (24). Recent evidence suggests that RAB1B, which has highly similar amino acid sequences with RAB1A, is required for trafficking the triglyceride-synthesizing enzyme DGAT2 to LDs in mammalian cells (33). *Drosophila* RAB1 is essential for targeting ER proteins to LDs (13). We hypothesized that MBOAT7 may act as a competitive inhibitor of RAB1A. Specifically, we hypothesized that MBOAT7 could restrict LD expansion in *CDS2* KO cells by competing with DGAT2 or GPAT4 for RAB1A binding, thereby interfering with their LD targeting. In line with this hypothesis, co-immunoprecipitation assays confirmed that both Flag-DGAT2 and Flag- GPAT4 interact with GFP-tagged RAB1A (Figure 4C and D). Interestingly, although MBOAT7 overexpression did not affect the binding between GPAT4 and RAB1A (Figure 4E), it significantly reduced the interaction between Flag-DGAT2 and GFP-RAB1A in *CDS2* KO cells (Figure 4F). When the interaction between MBOAT7 and RAB1A is disrupted, the binding between Flag-DGAT2 and GFP-RAB1A is retained (Figure 4F). Conversely, knockdown of MBOAT7 enhanced the binding between Flag-DGAT2 and GFP-RAB1A in *CDS2* KO cells (Figure 4G). To determine whether this regulated interaction translates to functional consequences in cells, confocal images showed that EGFP-DGAT2 was targeted to LDs in *CDS2* KO cells, but overexpression of MBOAT7-mCherry blocked this targeting (Figure 4H). These results collectively demonstrate that MBOAT7 specifically modulates the DGAT2-RAB1A interaction in a dose-dependent manner.

In response to OA treatment, triglyceride synthesis enzymes like DGAT2 or GPAT4 are more likely to target to LDs to promote LD growth (34). We also tested whether OA treatment affects the interaction of these enzymes with RAB1. OA treatment enhanced DGAT2 binding to RAB1A but did not affect GPAT4 binding to RAB1A (Figure 4I and Figure S2E). Interestingly, MBOAT7 binding to RAB1A increased after OA treatment in *CDS2* KO cells (Figure 4J). Taken together, our data suggest that CDS2 retains MBOAT7 at the ER. When CDS2 is absent, MBOAT7 binds to RAB1A and localizes to the ER-LD contacts. Furthermore, MBOAT7 reduces the size of LDs by reducing the targeting of DGAT2 to LDs.

### MBOAT7 and CDS2 coordinately regulate hepatic cholesterol excretion and fatty acid synthesis

The above results demonstrated that MBOAT7 and CDS2 regulate LD size in cells. Next, we examined how MBOAT7 and CDS2 functionally interact and regulate LD size *in vivo*. To define the coordinated roles of these enzymes in hepatic lipid metabolism, we performed genetic knockdown and overexpression studies in mice. We specifically single and double knocked down these enzymes in the liver, by injecting C57BL/6 mice with AAV-TBG-sh*Mboat7*, AAV-TBG-sh*Cds2* or AAV-TBG-sh*Mboat7*+AAV-TBG-sh*Cds2* (Figure 5A). Consistent with previous report (35), knockdown of *Mboat7* in liver did not affect plasma lipid content but increased hepatic triglyceride and cholesterol content (Figure 5B-E). Similarly, knockdown of *Cds2* did not affect plasma triglyceride content, but increased plasma and hepatic cholesterol and hepatic triglyceride content (Figure 5B-E). Interestingly, knockdown of both *Mboat7* and *Cds2* did not exert additive effects on the increased lipid levels, suggesting that they likely operate within the same regulatory network. The triglyceride and cholesterol contents were compatible to single knockdowns (Figure 5B-E). Consistent with the above results, H&E staining showed that knockdown of *Mboat7*, *Cds2*, and *Mboat7* + *Cds2* increased hepatic steatosis (Figure 5F-G).

**Figure 5.**
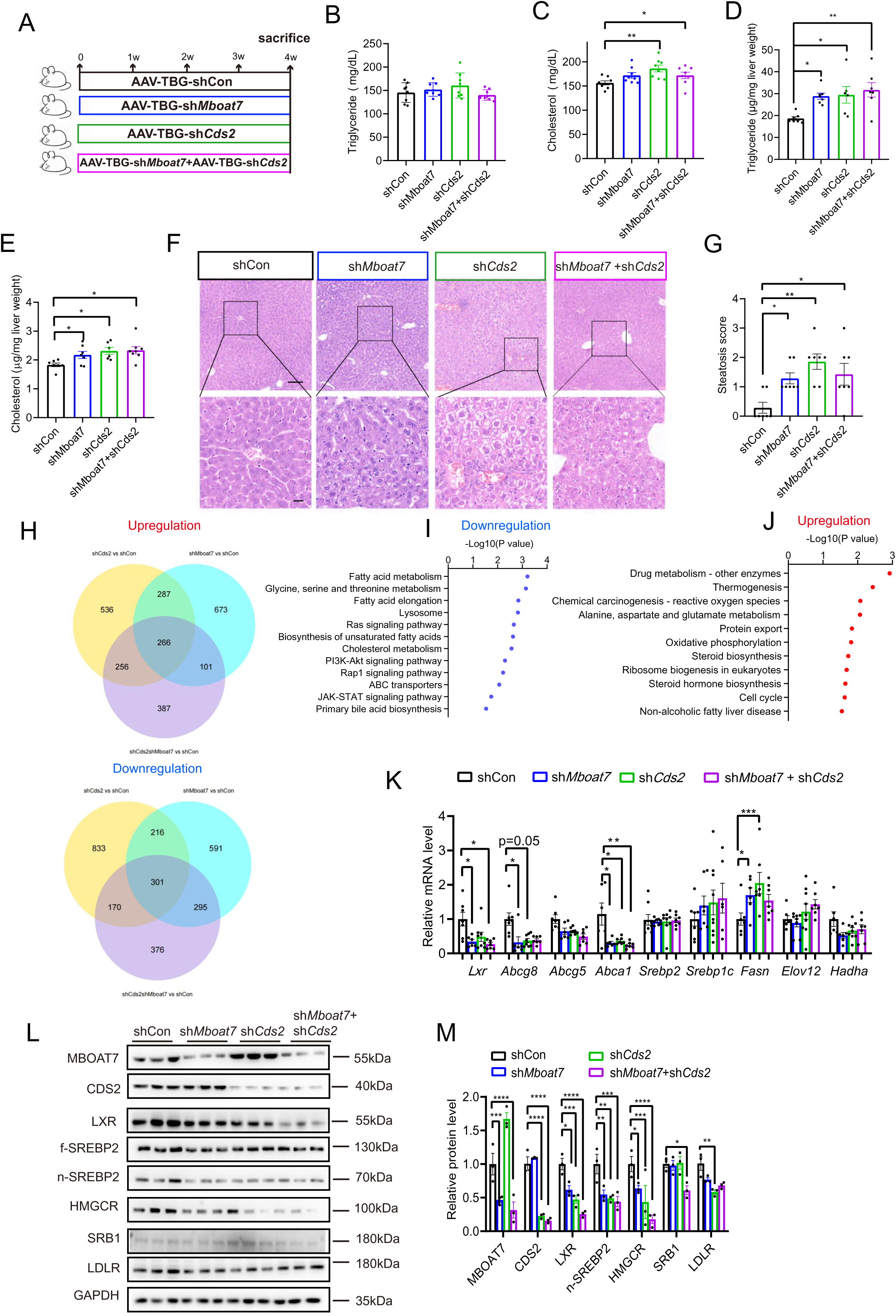
MBOAT7 and CDS2 coordinately regulate hepatic lipid metabolism. (A-E) Procedure for liver-specific knockdown of *Mboat7* and/or *Cds2* in mice. Mice were injected with AAV-TBG-shCon, AAV-TBG-sh*Mboat7*, AAV-TBG-sh*Cds2*, or AAV-TBG-sh*Mboat7* + AAV-TBG-sh*Cds2* and fed a chow diet for 4 weeks before analysis (A). Plasma triglyceride (B) and cholesterol (C), as well as hepatic triglyceride (D) and cholesterol (E) levels were measured in the mice in (A). N=6-7 mice per group. (F) H&E staining of liver sections from mice in (A). Scale bar represents 50 μm in main images and 10 μm in enlarged images. (G) Hepatic steatosis score based on liver sections of mice in (A). N=6-7 mice per group. (H) RNAseq was performed on liver samples from the mice in A. The numbers of genes up- or down-regulated in sh*Mboat7* vs sh*Con*, sh*Cds2* vs sh*Con* and sh*Mboat7* + sh*Cds2* vs sh*Con* are shown. (I) KEGG analysis results showing the pathways related to the 301 common downregulated genes in (A). (J) KEGG analysis results showing the pathways related to the 266 common upregulated genes in (A). (K) qRT-PCR analysis of the mRNA levels of genes that regulate cholesterol and fatty acid metabolism in the liver. N=5-8 mice per group. (L) Western blot analysis of liver lysates. N=3 mice per group. (M) Quantification of protein levels in (D). * p < 0.05, ** p < 0.01, *** p < 0.001, **** p < 0.0001.

To examine the pathways regulated by MBOAT7 and CDS2, we performed RNA sequencing (RNAseq) analysis on liver from the sh*Mboat7*, sh*Cds2*, and sh*Mboat7* + sh*Cds2* groups. When we compared the results, we noticed an extensive overlap between the three datasets: 266 genes were significantly upregulated and 301 genes were significantly downregulated in all three groups (Figure 5H). Kyoto Encyclopedia of Genes and Genomes (KEGG) analysis revealed significant changes in cholesterol and fatty acid metabolism pathways among the genes with downregulated expression in all three groups (Figure 5I). For the genes with upregulated expression in all three groups, KEGG analysis revealed significant alterations in steroid biosynthesis and non-alcoholic fatty liver pathways (Figure 5J).

LXR, a key regulator of cholesterol efflux, controls the transcription levels of genes such as *Abca1* and *Abcg5/8*. Our qPCR analysis verified that genes involved in cholesterol excretion (*Lxr, Abcg8* and *Abca1*) were downregulated in liver (Figure 5K), indicating that reduced expression of *Mboat7* or *Cds2* impairs cholesterol excretion. Knockdown of *Mboat7* and *Cds2* increased the expression of *Fasn*, which regulates fatty acid biosynthesis (Figure 5K). Accordingly, LXR protein levels were reduced in the liver of sh*Mboat7*, sh*Cds2*, and sh*Mboat7*+sh*Cds2* mice (Figure 5L-M). Nuclear SREBP2 and HMGCR levels were diminished, likely due to feedback inhibition of cholesterol biosynthesis by increased cellular cholesterol levels (Figure 5L-M). Together, these data suggests that MBOAT7 and CDS2 function in a coordinated way to orchestrate hepatic lipid metabolism.

### MBOAT7 regulates LD size and lipolysis in the absence of CDS2

Next, we examined whether MBOAT7 overexpression is able to reduce *Cds2* depletion-caused steatosis *in vivo*. *Cds2^f/f^* and *Cds2^f/f^;AlbCre* mice were injected with AAV-TBG-Con or AAV-TBG-*Mboat7* to specifically express *Mboat7* in liver (Figure 6A). Cell fractionation assays revealed that overexpression of *Mboat7* reduced the protein level of DGAT2 in the LD fraction from the livers of *Cds2^f/f^;AlbCre* mice, while increasing the protein level of ATGL (also called PNPLA2) (Figure 6B), a rate-limiting enzyme for lipolysis. This observation is consistent with our findings in cultured cells (Figure 4H), where MBOAT7 disrupted DGAT2 localization to LDs. The concurrent upregulation of ATGL further suggests that MBOAT7 expression may actively shift lipid metabolic balance toward lipolysis. To test if MBOAT7 regulates lipolysis, we used isoprenaline (ISO) to stimulate lipolysis. The results showed that overexpression of MBOAT7 increased free fatty acid (FFA) release in both DMSO and ISO-treated *Cds2*-deficient primary hepatocytes (Figure 6C).

**Figure 6.**
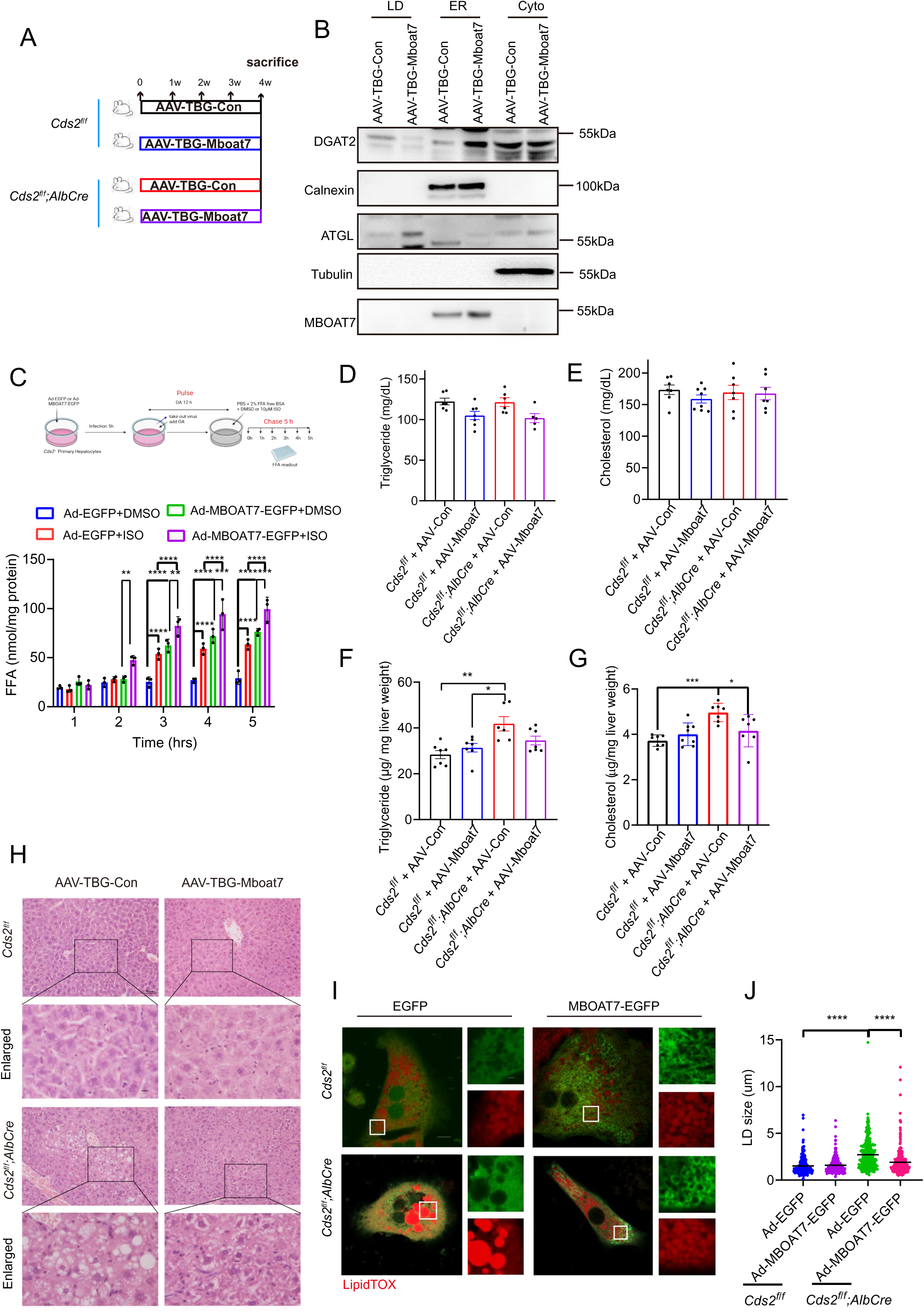
MBOAT7 regulates LD size and lipolysis. (A) Schematic showing the procedure for hepatic overexpression of MBOAT7 in the liver of mice with and without hepatic depletion of *Cds2*. *Cds2^f/f^* and *Cds2^f/f^;AlbCre* mice were injected with AAV-TBG-Con or AAV-TBG-*Mboat7* and fed a chow diet for 4 weeks before analysis. (B) Western blot analysis of LD, ER and cytosol fractions from the liver of *Cds2^f/f^;AlbCre* mice that are injected with AAV-TBG-Con or AAV-TBG-*Mboat7*. (C) Primary hepatocytes from *Cds2^f/f^* and *Cds2^f/f^;AlbCre* mice were infected with Ad-EGFP or Ad-MBOAT7-EGFP, then treated with 10 μM ISO or DMSO (control) for the indicated times. Free fatty acids in the medium were analyzed. (D and E) Plasma triglyceride (D) and cholesterol (E) levels were analyzed in the mice from (B). (F and G) Hepatic triglyceride (F) and cholesterol (G) levels were analyzed in the mice from (A). N=7-8 mice per group. (H) H&E staining of liver sections from *Cds2^f/f^* and *Cds2^f/f^;AlbCre* mice. Scale bar represents 50 μm in main images and 10 μm in enlarged images. (I) Primary hepatocytes isolated from *Cds2^f/f^* and *Cds2^f/f^;AlbCre* mice were infected with Ad-EGFP or Ad-MBOAT7-EGFP and treated with 100 μM OA for 12 hours. LDs were stained with LipdTOX before confocal imaging. Scale bar represents 10 μm in main images and 2 μm in enlarged images. (J) Quantification of LD sizes in (I). * p < 0.05, ** p < 0.01, *** p < 0.001, **** p < 0.0001.

Hepatic overexpression of *Mboat7* did not affect plasma lipids in either control or *Cds2^f/f^;AlbCre* mice (Figure 6D-E). Hepatic overexpression of *Mboat7* did not change hepatic triglyceride and cholesterol levels in *Cds2^f/f^* mice, but it reduced hepatic triglyceride and cholesterol levels in *Cds2^f/f^;AlbCre* mice (Figure 6F-G). H&E staining showed that liver sections from *Cds2^f/f^;AlbCr*e mice contained “supersized” LDs, while overexpression of *Mboat7* reduced the LD size in *Cds2^f/f^;AlbCre* mice, resulting in multiple small-sized LDs in hepatocytes (Figure 6H). This reduction in LD size was also confirmed in *Cds2*-deficient primary hepatocytes (Figure 6I-J). Overall, our data suggest that MBOAT7 controls LD size and lipolysis by inhibiting DGAT2 targeting to LDs and promoting ATGL targeting to LDs in both *CDS2*-deficient cell lines and primary hepatocytes.

### MBOAT7 and CDS2 regulate cholesterol excretion through the AMPK pathway

Given that knockdown of *Cds2* and/or *Mboat7* increased hepatic cholesterol levels and overexpression of *Mboat7* reduced hepatic cholesterol levels in *Cds2^f/f^;AlbCr*e mice, we investigated the mechanism by which CDS2 and MBOAT7 regulate hepatic cholesterol metabolism. Our results showed that knockdown of *Mboat7* reduced the expression levels of genes in the liver X receptor (LXR) pathway, suggesting impaired cholesterol excretion (Figure 5). Heatmap analysis further confirmed the downregulation of genes (*Abcg5* and *Abcg8*) involved in cholesterol excretion (Figure 7A). Activation of the AMPK pathway is known to inhibit SREBP and enhance LXR protein levels. AMPK signaling was downregulated in the liver of sh*Mboat7*, sh*Cds2*, or sh*Mboat7* + sh*Cds2* mice, as indicated by heatmap analysis and western blotting (Figure 7A and Figure S3A-B). In contrast, overexpression of *Mboat7* increased p-AMPK and p-ACC levels in *Cds2^f/f^;AlbCre* mice, suggesting that AMPK pathway activity was enhanced (Figure 7B and Figure S3C). This was accompanied by reduced nuclear SREBP2 levels and increased expressions of LXR and its target ABCA1 in *Cds2^f/f^;AlbCre* mice (Figure 7B). Accordingly, mRNA analysis revealed that downstream targets of LXR (*Abca1*, *Abcg5* and *Abcg8*) were upregulated in the liver of *Mboat7*-overexpressing *Cds2^f/f^;AlbCre* mice (Figure 7C).

**Figure 7.**
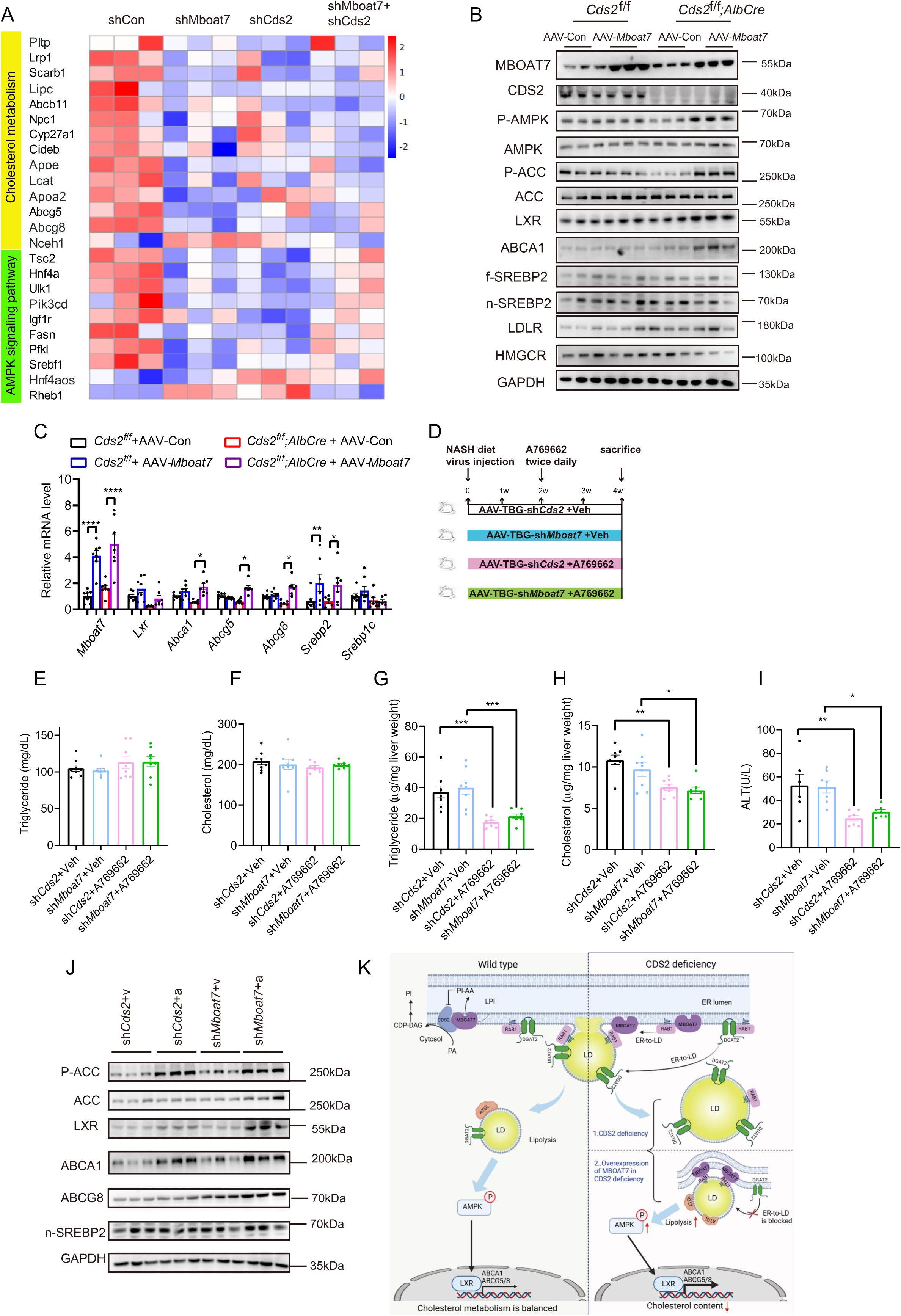
MBOAT7 and CDS2 regulate cholesterol metabolism through the AMPK pathway. (A) Heatmap analysis shows alterations of expression levels of genes involved in cholesterol metabolism and the AMPK pathway in the liver of 8-week-old mice injected with AAV-TBG-sh*Con*, AAV-TBG-sh*Mboat7*, AAV-TBG-sh*Cds2*, and AAV-TBG-sh*Mboat7* + AAV-TBG-sh*Cds2*. (B) Western blot of *Cds2^f/f^* and *Cds2^f/f^;AlbCre* mice injected with AAV-TBG-Con or AAV-TBG-*Mboat7*, and fed a chow diet for 4 weeks before analysis. (C) qRT-PCR analysis of genes involved in cholesterol and fatty acid metabolism. (D) Schematic showing the procedure for sh*Cds2* and sh*Mboat7* mice with or without A769662 treatment. 8-week-old C57BL6/J mice were injected with either AAV-TBG-sh*Cds2* or AAV-TBG-sh*Mboat7* and concurrently fed a NASH-inducing GAN diet for 4 weeks. During the final 2 weeks of this period, mice received intraperitoneal injections of the AMPK agonist A769662 (30 mg/kg, twice daily). (E-H) Plasma triglyceride (E) and cholesterol (F), hepatic triglyceride (G) and cholesterol (H), plasma ALT levels (I) were shown. (J) Protein levels of key components in the LXR signaling pathway. (K) A model illustrating how MBOAT7 and CDS2 regulate LD dynamics and cholesterol metabolism in hepatocytes. CDS2 and MBOAT7 physically interact in the ER membrane and both regulate PI biosynthesis. MBOAT7 catalyzes the generation of PI-AA, which inhibits CDS2 activity. In the absence of CDS2, MBOAT7 relocates to the ER-LD contacts via binding to RAB1. CDS2 deficiency causes increased LD size accompanied by increased recruitment of DGAT2 to LDs, whereas overexpression of MBOAT7 in CDS2-deficient conditions reduces DGAT2 targeting to LDs, increases ATGL recruitment to the LD surface, and reduces LD size. The increased lipolysis induced by MBOAT7 expression activates AMPK, enhancing LXR protein levels and expression of its downstream target genes, thus stimulating cholesterol metabolism. * p < 0.05, ** p < 0.01, *** p < 0.001, **** p < 0.0001,

We then investigated whether pharmacological activation of AMPK could reverse the MASLD phenotype in *Cds2*- or *Mboat7*-deficient mice. 8-week-old mice were injected with either AAV-TBG-sh*Cds2* or AAV-TBG-sh*Mboat7* and concurrently fed a NASH-inducing GAN diet for 4 weeks. During the final 2 weeks of this period, mice received intraperitoneal injections of the AMPK agonist A769662 (30 mg/kg, twice daily; Figure 7D). Following A769662 treatment, body weight was reduced in both *Cds2*- and *Mboat7*-deficient mice (Figure S3D). Although plasma triglyceride and cholesterol levels remained unchanged, hepatic levels of triglyceride and cholesterol were significantly decreased (Figure 7E-H). Additionally, A769662 treatment lowered plasma ALT levels, indicating reduced liver inflammation (Figure 7I). H&E and Oil-Red-O staining confirmed a marked reduction in hepatic lipid accumulation in A769662-treated mice compared to vehicle controls (Figure S3E). To assess downstream signaling, we examined the AMPK-LXR axis. A769662 treatment increased protein levels of p-ACC, LXR and ABCA1 in sh*Cds2* and sh*Mboat7* livers (Figure 7J and Figure S3F). Together, these findings suggest that MBOAT7 and CDS2 regulate cholesterol excretion through activation of the AMPK-LXR axis.

## Discussion

Here, we elucidate that spatial localization of MBOAT7 to ER-LD contact sites-orchestrated by CDS2-represents a fundamental cellular adaptation to lipid overload. This redistribution is critical for its role in regulating LD dynamics. At the ER, CDS2 binds to MBOAT7, but in the absence of CDS2, MBOAT7 binds to RAB1A and locates to the ER-LD contacts. There, by sequestering RAB1A, MBOAT7 competitively disrupts the RAB1A-DGAT2 interaction, thereby inhibiting DGAT2-mediated LD growth and promoting lipolysis (Figure 7K).

LDs play an essential role in adaptation to fluctuating metabolic states, through controlling their size, subcellular distribution and interaction with other organelles (36). Under lipid overload, cells sequester free fatty acid as triglyceride within the core of LDs to prevent lipotoxicity and oxidative stress. Concurrently, they activate counterregulatory mechanisms to prevent excessive LD accumulation (4). Phospholipid metabolic enzymes, which supply lipids for LD formation, are actively involved in this process. Our study identifies MBOAT7 and CDS2 as key coordinators of LD growth. In MASLD mouse livers, MBOAT7 is more concentrated on LDs compared to normal controls, suggesting a compensatory mechanism to restrain steatosis-driven LD expansion. This represents a protective response in MASLD pathogenesis, whereby the redistribution of an ER protein to ER-LD contacts serves to limit LD growth under lipid overload.

ER-LD contact is a critical metabolic hub that maintain organelle function and metabolic homeostasis (8, 12, 37). Although lipid metabolic enzymes are present at these sites, their precise roles are still being defined. Emerging evidence underscores the importance of ER-LD contacts as central platforms for lipid, particularly phospholipid metabolism (38, 39). For instance, LPCAT3, a member of the MBOAT family, localizes at the ER-LD contacts and remodels the acyl-chain composition of phosphatidylethanolamine (PE) to influence LD growth (38). In contrast, MBOAT7 regulates LD size independent of its enzymatic activity, highlighting the diverse strategies employed at these contact sites. The broader physiological significance of ER-LD contacts in coordinating lipid metabolism is exemplified In *Cds2* deficient hepatocytes. The compensatory redistribution of MBOAT7 to ER-LD contacts stimulates ATGL-mediated lipolysis, reducing LD size, elevating cytosolic free fatty acids, and promoting cholesterol excretion. Thus, ER-LD contacts function not merely as passive tethers for protein trafficking, but dynamic regulatory nodes that integrate phospholipid remodeling with neutral lipid turnover, ensuring metabolic adaptation and homeostasis.

RAB1 is a central player of ER-to-Golgi transport, also plays a central role in recruiting multiple lipid metabolizing enzymes to LDs, thereby fine-tuning LD dynamics and lipid homeostasis (13, 33). Here we found that RAB1A interacts with MBOAT7 and promotes its redistribution to LD surface. This interaction competes DGAT2-mediated LD growth, shifting the metabolic balance toward lipolysis. Importantly, this competitive recruitment only becomes evident under CDS2 deficiency, suggesting that RAB1 exhibits a stronger binding affinity for DGAT2 than for MBOAT7. Given that RAB1A and RAB1B share 92% amino acid identity and display functional redundancy (40), our findings consolidate a model in which RAB1 serves as a multimodal adaptor that dynamically orchestrate the LD localization of key enzymes involved in both neutral lipid and phospholipid metabolism.

In summary, our findings reveal a previously unrecognized axis of inter-organelle communication and metabolic control, orchestrated by MBOAT7, CDS2, and RAB1A that adaptively responds to lipid overload. We provide a new mechanistic link between organelle membrane dynamics and metabolic homeostasis in MASLD pathogenesis.

## Material and Methods

### Mice

*Cds2^f/f^* and *Cds2^f/f^;AlbCre* mice were generated as described (26). C57BL/6J mice were purchased from GemPharmatech Co., Ltd. For liver-specific gene knockdowns, C57BL/6J mice were injected with AAV-TBG-sh*Con*, AAV-TBG-sh*Mboat7*, AAV-TBG-sh*Cds2* intravenously (i.v) with 3×10^11^ genomic copies of AAVs. The target sequence for *Mboat7* is 5’-CCTGTTGGCTTCCTCTTTAAG-3’. The target sequence for *Cds2* is 5’-GTCCGAAGCAAAGCTAGAT-3’. The control sequence is 5’-GAAGTCGTGAGAAGTAGAA-3’. For liver-specific overexpression of *Mboat7*, C57BL/6J mice were injected with AAV-TBG-*Con* or AAV-TBG-*Mboat7* intravenously (i.v) with 1×10^11^ genomic copies of AAVs. Mice were housed at room temperature (22°C) and fed a standard rodent chow diet (Beijing HFK Bioscience Co., Ltd #1032). Unless otherwise stated, 8-week-old male mice were used and all mice were fasted for 5-6 hours prior to euthanization. In some experiments, C57BL/6J mice were fed a chow or GAN diet (Research Diets Inc, D09100310) for 26 weeks. All animal experiments were approved by the Institutional Animal Care and Use Committee (IACUC) at the Institute of Genetics and Developmental Biology, Chinese Academy of Sciences (IGDB-CAS).

### Cell culture

The *CDS2* knockout HeLa cell line was generated using the CPISPR/Cas9 system according to published literature (41). The target sequence was 5’-TATTTACTGAGAATCCGCAA-3’. HepG2 cells (ATCC, HB-8065) and HeLa cells were cultured in DMEM with 10% FBS. In certain experiments, HeLa cells were transfected with siRNAs. The targeting sequence for *RAB1A* is 5’-CAGCATGAATCCCGAATAT-3’. The targeting sequence for *CDS2* is 5’-GGCCACCTTTGTCAATGTA-3’. Mouse primary hepatocytes were isolated from *Cds2^f/f^* and *Cds2^f/f^;AlbCre* mice and infected with Ad-*EGFP* or Ad-*MBOAT7-EGFP* with MOI=50. All cells were grown at 37°C in a 5% CO_2_ incubator.

### APEX-based proximity labeling

The cDNA encoding CDS2 was cloned into *pcDNA3APEX2-NES* vector, generating C-terminal fusion protein. Cells expressing the APEX2-CDS2 were incubated with 500µM biotin-phenol for 1 hour at 37⁰C, followed by a 1-minute labeling reaction initiated by the addition of 1 mM H_2_O_2_. The reaction was promptly quenched with a TBS solution (Thermo #28376), and the cells were harvested using RIPA buffer. Following cell lysis, biotinylated proteins were affinity-purified using streptavidin-coated beads. After washing, the captured proteins were subjected to on-bead tryptic digestion, and the resulting peptides were analyzed by liquid chromatography-tandem mass spectrometry (LC-MS/MS). Specific proximal interactors were distinguished from background by comparison with cells expressing APEX2 alone.

### Plasmids

Human *MBOAT7* cDNA was cloned into the *pcDNA3.1-EGFP* and *pCMV-C-mCherry* vectors. Human *CDS2* cDNA was cloned into the *pCMV-Tag2b* and *pCS2-HA* vectors. Human *RAB1A*, *DGAT2* and *GPAT4* cDNAs were cloned into both the *pEGFP-N1* and *pCMV-Tag2b* vectors.

### MBOAT7 activity assay

The activity of MBOAT7 was measured as described with minor modifications (28). Briefly, 100 μg of protein from liver was incubated in 200 μl of medium (10 mM Tris/HCl, pH 7.5, 150 mM NaCl, 1 mM EDTA) with 50 μM lysophosphatidylinositol (LPI), 25 μM arachidonoyl-CoA (20:4-CoA), and 5 μM (0.2 µCi) radiolabeled arachidonoyl[1-14C]CoA. The reaction was incubated for 15-30 minutes at 37 ⁰C and then stopped by adding 1 mL of chloroform:methanol (2:1, vol/vol) and 0.4 mL of 2% orthophosphoric acid. After centrifugation (3,500 g, 15 min, 4 ⁰C), the lower organic phase, containing the lipids, was collected and dried under a gentle nitrogen stream. Lipids were reconstituted in chloroform. Total lipids were extracted using the Bligh and Dyer method and separated by one-dimensional TLC on silica gel 60 plates using chloroform/methyl acetate:1-propanol:methanol:0.25% KCl (25:25:25:10:9, vol/vol). The radioactivity in the separated bands were quantified using a Typhoon FLA 9000 (GE Healthcare Japan). The corresponding areas of radiolabeled phospholipids on TLC plates were cut out, and the radiolabeled products were measured by liquid scintillation counting.

### Lipid analysis

Lipids were extracted based on the chloroform/methanol (2:1, v/v) lipid extraction method (3). In brief, 50 mg of liver tissue were extracted with 6 mL of chloroform/methanol (2:1, v/v), then vortexed and refrigerated overnight. 3 mL of 0.043% magnesium chloride was then added to separate the aqueous phase from the chloroform phase which contains the lipids. After separation of the two phases, the upper water/methanol phase was removed. A 0.5-mL aliquot of the bottom phase was combined with 1 mL of 0.5% Triton X-100 in chloroform and dried. Triglyceride and total cholesterol levels were measured using commercially available kits (BioSino, Beijing, China). Plasma free fatty acids (FFA) were measured using commercially available kits (Sigma-Alrich, MAK044).

### Immunoprecipitation

Cells were transfected with the indicated plasmids using Lipofectamine 3000 (ThermoFisher, L3000001). Proteins were extracted using RIPA lysis buffer, containing100 mM Tris Base, 1% deoxycholate, 0.1% SDS, 5 mM EDTA, 1% Triton-X-100, 150 mM NaCl, pH 8.0), supplemented with 100 mM PMSF, protease inhibitor cocktail (Roche, P8340) and 100 mM sodium vanadate. Lysates were incubated on ice for 15 min, then centrifuged at 12,000 rpm for 15 minutes at 4 °C. Supernatants were incubated overnight at 4 °C with anti-Flag M2 affinity gel (Sigma-Aldrich, A2220) or anti-GPF agarose beads (Lablead Biotech Co., Ltd). After four washes with lysis buffer, beads were denatured in 2X SDS sample buffer for 10 min at 100 °C for western blotting. For the pulldown assay, MBOAT7-His was pulled down using Ni-NTA agarose beads (Qiagen, 30210), then washed three times with PBS + PMSF + 1% Triton buffer. 10 μg of *Flag-CDS2* was expressed in 293T cells, immunoprecipitated using anti-Flag M2 affinity gel, and eluted with 3X Flag peptide. The eluted Flag-CDS2 (1.5 mL) was then incubated with the Ni-NTA agarose-bound MBOAT7-His for 2 hours. The immunoprecipitate was washed six times with PBS + PMSF + 1% Triton buffer and denatured with 2X SDS sample buffer for 10 min before western blotting.

### Confocal imaging

The images were acquired using a LSM 980 with Airyscan 2 (Zeiss) or a confocal microscope (Leica SP8). Images were analyzed by Image J.

### Subcellular fractionation

Subcellular fractionation of liver tissue was performed as described (42).

### Antibodies

Rabbit antibodies against CDS2 (1:500, 13175-1-AP), SREBP2 (1:1000, 28212-1-AP), LDLR (1:1000, 10785-1-AP), SR-B1 (1:1000, 21277-1-AP) and MBOAT7 (1:100, 83546-3-RR) were purchased from Proteintech. Rabbit antibody against MBOAT7 (1:1000, AV49811) was purchased from Sigma-Aldrich. Mouse antibody against Flag (1:1000, F1804) was purchased from Sigma-Aldrich. Rabbit antibody against Flag (1:2500, PM020) was purchased from MBL. Rabbit antibody against ATGL (1:1000, 2138) was purchased from Cell Signaling Technology. Mouse antibody against LXR (1:1000, ab41902) was purchased from Abcam. Rabbit antibodies against DGAT2 (1:1000, ab237613) and HMGCR (1:1000, ab174830) were purchased from Abcam.

### Statistics

Results with error bars are mean±SEM. Comparisons between 2 groups were performed using the two-tailed Student’s t-test. ANOVA was used for comparisons between more than 2 groups.

## Acknowledgments

We thank Xixian Li from the Center for Biological Imaging at the Institute of Biophysics, CAS, for analyzing images. This work was supported by grants 2024YFA1306101, 2023YFA1800500 from National Key R&D Program of China and 92354301, 32230044, 32371230 and 32321004 from the National Natural Science Foundation of China, and BJ-2024-235 from National High Level Hospital Clinical Research Funding.

Figure 3A, Figure 6C and Figure 7K were created with BioRender (https://BioRender.com/ *k88h902*, https://BioRender.com/s73×778*, and* https://BioRender.com/e57c295).

## Author Contributions

J Xu and X Huang designed the experiments. J Xu, J Zhang, W Wang and H Su performed the experiments. J Xu, J Zhang, S Chen and M Ding analyzed the data. J Xu and X Huang wrote the manuscript.

## References

1. L. Xiang et al., A multi-omic landscape of steatosis-to-NASH progression. Life Metab 1, 242–257 (2022).

2. T. C. Walther, J. Chung, R. V. Farese, Jr., Lipid Droplet Biogenesis. Annu Rev Cell Dev Biol 33, 491–510 (2017).

3. E. Scorletti, R. M. Carr, A new perspective on NAFLD: Focusing on lipid droplets. J Hepatol 76, 934–945 (2022).

4. J. L. Dempsey, G. N. Ioannou, R. M. Carr, Mechanisms of Lipid Droplet Accumulation in Steatotic Liver Diseases. Semin Liver Dis 43, 367–382 (2023).

5. N. L. Gluchowski, M. Becuwe, T. C. Walther, R. V. Farese, Jr., Lipid droplets and liver disease: from basic biology to clinical implications. Nat Rev Gastroenterol Hepatol 14, 343–355 (2017).

6. M. B. Schott et al., Lipid droplet size directs lipolysis and lipophagy catabolism in hepatocytes. J Cell Biol 218, 3320–3335 (2019).

7. G. F. Grabner, H. Xie, M. Schweiger, R. Zechner, Lipolysis: cellular mechanisms for lipid mobilization from fat stores. Nat Metab 3, 1445–1465 (2021).

8. K. Qian et al., CLSTN3beta enforces adipocyte multilocularity to facilitate lipid utilization. Nature 613, 160–168 (2023).

9. G. E. Miner et al., PLIN5 interacts with FATP4 at membrane contact sites to promote lipid droplet-to-mitochondria fatty acid transport. Dev Cell 58, 1250–1265 e1256 (2023).

10. A. Kumar, S. Yadav, V. Choudhary, The evolving landscape of ER-LD contact sites. Front Cell Dev Biol 12, 1483902 (2024).

11. M. J. Olarte et al., Determinants of Endoplasmic Reticulum-to-Lipid Droplet Protein Targeting. Dev Cell 54, 471–487 (2020).

12. D. Li et al., The ER-Localized Protein DFCP1 Modulates ER-Lipid Droplet Contact Formation. Cell Rep 27, 343–358 e345 (2019).

13. J. Song et al., Identification of two pathways mediating protein targeting from ER to lipid droplets. Nat Cell Biol 24, 1364–1377 (2022).

14. A. Caddeo, R. Spagnuolo, S. Maurotti, MBOAT7 in liver and extrahepatic diseases. Liver Int 43, 2351–2364 (2023).

15. V. R. Thangapandi et al., Loss of hepatic Mboat7 leads to liver fibrosis. Gut 70, 940–950 (2021).

16. V. Varadharajan, W. J. Massey, J. M. Brown, Membrane-bound O-acyltransferase 7 (MBOAT7)-driven phosphatidylinositol remodeling in advanced liver disease. J Lipid Res 63, 100234 (2022).

17. J. Alharthi et al., A metabolic associated fatty liver disease risk variant in MBOAT7 regulates toll like receptor induced outcomes. Nat Commun 13, 7430 (2022).

18. M. Xia, P. Chandrasekaran, S. Rong, X. Fu, M. A. Mitsche, Hepatic deletion of Mboat7 (LPIAT1) causes activation of SREBP-1c and fatty liver. J Lipid Res 62, 100031 (2021).

19. Y. Liu, W. Wang, G. Shui, X. Huang, CDP-diacylglycerol synthetase coordinates cell growth and fat storage through phosphatidylinositol metabolism and the insulin pathway. PLoS Genet 10, e1004172 (2014).

20. N. J. Blunsom, S. Cockcroft, CDP-Diacylglycerol Synthases (CDS): Gateway to Phosphatidylinositol and Cardiolipin Synthesis. Front Cell Dev Biol 8, 63 (2020).

21. K. D’Souza, Y. J. Kim, T. Balla, R. M. Epand, Distinct properties of the two isoforms of CDP-diacylglycerol synthase. Biochemistry 53, 7358–7367 (2014).

22. M. P. Moore et al., Low MBOAT7 expression, a genetic risk for MASH, promotes a profibrotic pathway involving hepatocyte TAZ upregulation. Hepatology 81, 576–590 (2024).

23. Y. Tanaka et al., LPIAT1/MBOAT7 depletion increases triglyceride synthesis fueled by high phosphatidylinositol turnover. Gut 70, 180–193 (2021).

24. Y. Xu et al., CDP-DAG synthase 1 and 2 regulate lipid droplet growth through distinct mechanisms. J Biol Chem 294, 16740–16755 (2019).

25. Y. Qi et al., CDP-diacylglycerol synthases regulate the growth of lipid droplets and adipocyte development. J Lipid Res 57, 767–780 (2016).

26. J. Xu et al., Hepatic CDP-diacylglycerol synthase 2 deficiency causes mitochondrial dysfunction and promotes rapid progression of NASH and fibrosis. Sci Bull (Beijing) 67, 299–314 (2022).

27. J. A. Olzmann, P. Carvalho, Dynamics and functions of lipid droplets. Nat Rev Mol Cell Biol 20, 137–155 (2019).

28. K. Wang et al., The structure of phosphatidylinositol remodeling MBOAT7 reveals its catalytic mechanism and enables inhibitor identification. Nat Commun 14, 3533 (2023).

29. R. Dhiman, S. Caesar, A. R. Thiam, B. Schrul, Mechanisms of protein targeting to lipid droplets: A unified cell biological and biophysical perspective. Semin Cell Dev Biol 108, 4–13 (2020).

30. C. I. Pataki et al., Proteomic analysis of monolayer-integrated proteins on lipid droplets identifies amphipathic interfacial alpha-helical membrane anchors. Proc Natl Acad Sci U S A 115, E8172–E8180 (2018).

31. K. Bersuker et al., A Proximity Labeling Strategy Provides Insights into the Composition and Dynamics of Lipid Droplet Proteomes. Dev Cell 44, 97–112 e117 (2018).

32. N. Mejhert et al., Partitioning of MLX-Family Transcription Factors to Lipid Droplets Regulates Metabolic Gene Expression. Mol Cell 77, 1251–1264 e1259 (2020).

33. Y. Malis et al., Rab1b facilitates lipid droplet growth by ER-to-lipid droplet targeting of DGAT2. Sci Adv 10, eade7753 (2024).

34. F. Wilfling et al., Triacylglycerol synthesis enzymes mediate lipid droplet growth by relocalizing from the ER to lipid droplets. Dev Cell 24, 384–399 (2013).

35. M. Meroni et al., Mboat7 down-regulation by hyper-insulinemia induces fat accumulation in hepatocytes. EBioMedicine 52, 102658 (2020).

36. R. W. Klemm, P. Carvalho, Lipid Droplets Big and Small: Basic Mechanisms That Make Them All. Annu Rev Cell Dev Biol 40, 143–168 (2024).

37. D. Xu et al., Rab18 promotes lipid droplet (LD) growth by tethering the ER to LDs through SNARE and NRZ interactions. J Cell Biol 217, 975–995 (2018).

38. M. J. Tol et al., Dietary control of peripheral adipose storage capacity through membrane lipid remodelling. Nat Metab 7, 1424–1442 (2025).

39. Q. Li, X. Zhou, X. Zhang, C. Zhang, S. O. Zhang, Nuclear receptor signaling regulates compartmentalized phosphatidylcholine remodeling to facilitate thermosensitive lipid droplet fusion. Nat Commun 16, 3955 (2025).

40. X. Z. Yang et al., Rab1 in cell signaling, cancer and other diseases. Oncogene 35, 5699–5704 (2016).

41. F. A. Ran et al., Genome engineering using the CRISPR-Cas9 system. Nat Protoc 8, 2281–2308 (2013).

42. Y. Ding et al., Isolating lipid droplets from multiple species. Nat Protoc 8, 43–51 (2013).

