## Supplemental Figure s1-3 for "A partner switch from CDS2 to RAB1A redirects MBOAT7 to inhibit DGAT2-mediated lipid droplet growth"

A

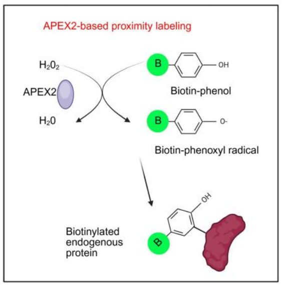


B

| Accession | Gene names | Function |
|---|---|---|
| O95674 | CDS2 | Catalyzes the conversion of phosphatidic acid (PA) to CDP-diacylglycerol (CDP-DAG) |
| P27824-3 | CALX | Calcium-binding protein that interacts with newly synthesized glycoproteins in the endoplasmic reticulum |
| Q92973-2 | TNPO1 | Functions in nuclear protein import as nuclear transport receptor |
| P55060-4 | CSE1L | Mediates importin-alpha re-export from the nucleus to the cytoplasm after import substrates (cargos) have been released into the nucleoplasm |
| Q14152-2 | EIF3A | NA-binding component of the eukaryotic translation initiation factor 3 (eIF-3) complex |
| P49368 | TCPG | A molecular chaperone complex that assists the folding of proteins upon ATP hydrolysis |
| O60610-2 | DIAP1 | Actin-binding |
| P22392-2 | NDKB | Major role in the synthesis of nucleoside triphosphates other than ATP |
| P52272-2 | HNRNPM | Ribonucleoprotein; RNA-binding |
| Q15293 | RCN1 | Regulation of Insulin-like Growth Factor (IGF) transport and uptake by Insulin-like Growth Factor Binding Proteins |
| Q14527-2 | HLTF | Has both helicase and E3 ubiquitin ligase activities |
| P33897 | ABCD1 | ATP-dependent transporter of the ATP-binding cassette (ABC) family involved in the transport of very long chain fatty acid (VLCFA)-CoA from the cytosol to the peroxisome lumen |
| Q93008-1 | USP9X | Deubiquitinase involved both in the processing of ubiquitin precursors and of ubiquitinated proteins |
| Q969M3 | YIPF5 | Plays a role in transport between endoplasmic reticulum and Golgi |
| P61224-2 | RAP1B | possesses intrinsic GTPase activity |
| P42677 | RS27 | Component of the small ribosomal subunit |
| O43169 | CYB5B | Cytochrome b5 is a membrane-bound hemoprotein functioning as an electron carrier for several membrane-bound oxygenases |
| Q13619-2 | CUL4A | Core component of multiple cullin-RING-based E3 ubiquitin-protein ligase complexes which mediate the ubiquitination of target proteins |
| Q5SRE5-2 | NUP188 | Function as a component of the nuclear pore complex (NPC). |
| Q8NBS9-2 | TXND5 | Possesses thioredoxin activity. Has been shown to reduce insulin disulfide bonds. |
| Q96N66-3 | MBOAT7 | Mediates the conversion of lysophosphatidylinositol (1-acylglycerophosphatidylinositol or LPI) into phosphatidylinositol (1,2-diacyl-sn-glycero-3-phosphoinositol or PI) (LPIAT activity) |
| Q01844-2 | EWSR1 | Function as a transcriptional repressor. |
| Q6UWP8 | SBSN | Function not clear |
| P25311 | AZGP1 | Stimulates lipid degradation in adipocytes and causes the extensive fat losses associated with some advanced cancers. May bind polyunsaturated fatty acids. |

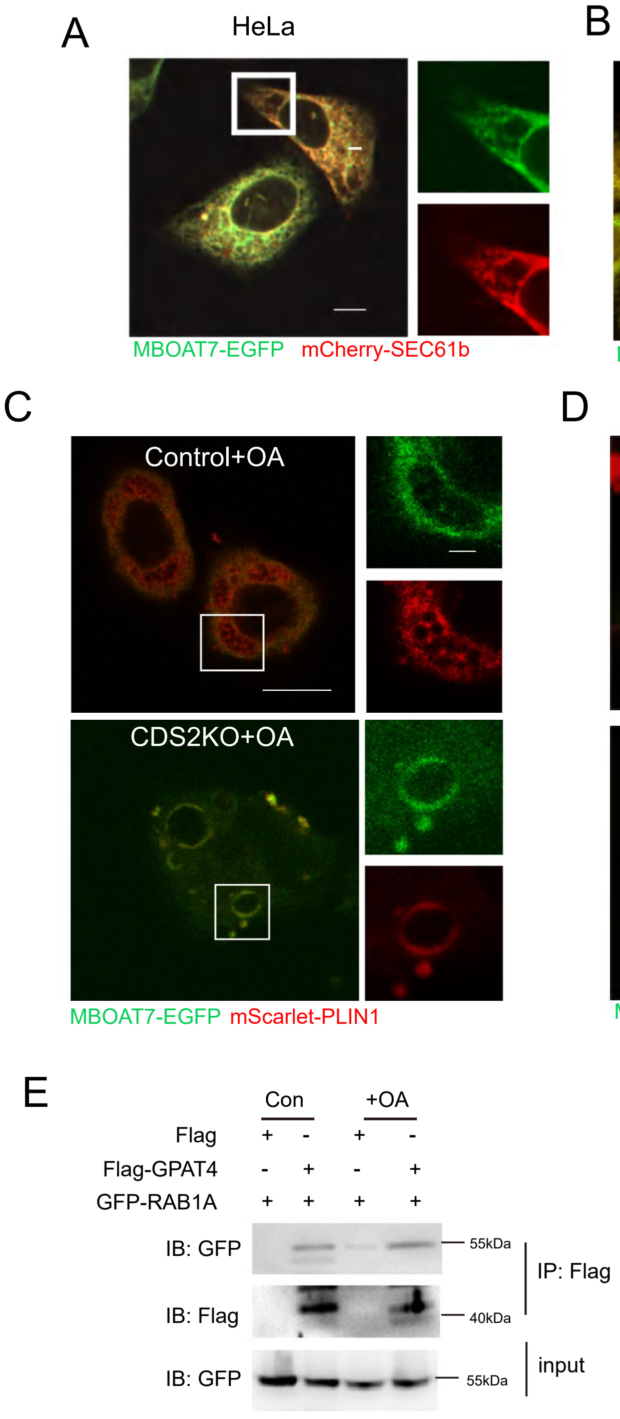
A
HeLa
MBOAT7-EGFP
mCherry-SEC61b
B
CDS2KO
MBOAT7-EGFP
mCherry-SEC61b
C
Control+OA
CDS2KO+OA
MBOAT7-EGFP
mScarlet-PLIN1
D
HepG2 + OA
NC
siCDS2
MBOAT7-EGFP
LipidTOX
E
Con
+OA
Flag + - + -
Flag-GPAT4 - + - +
GFP-RAB1A + + + +
IB: GFP
IB: Flag
IB: GFP
55kDa
40kDa
55kDa
IP: Flag
input

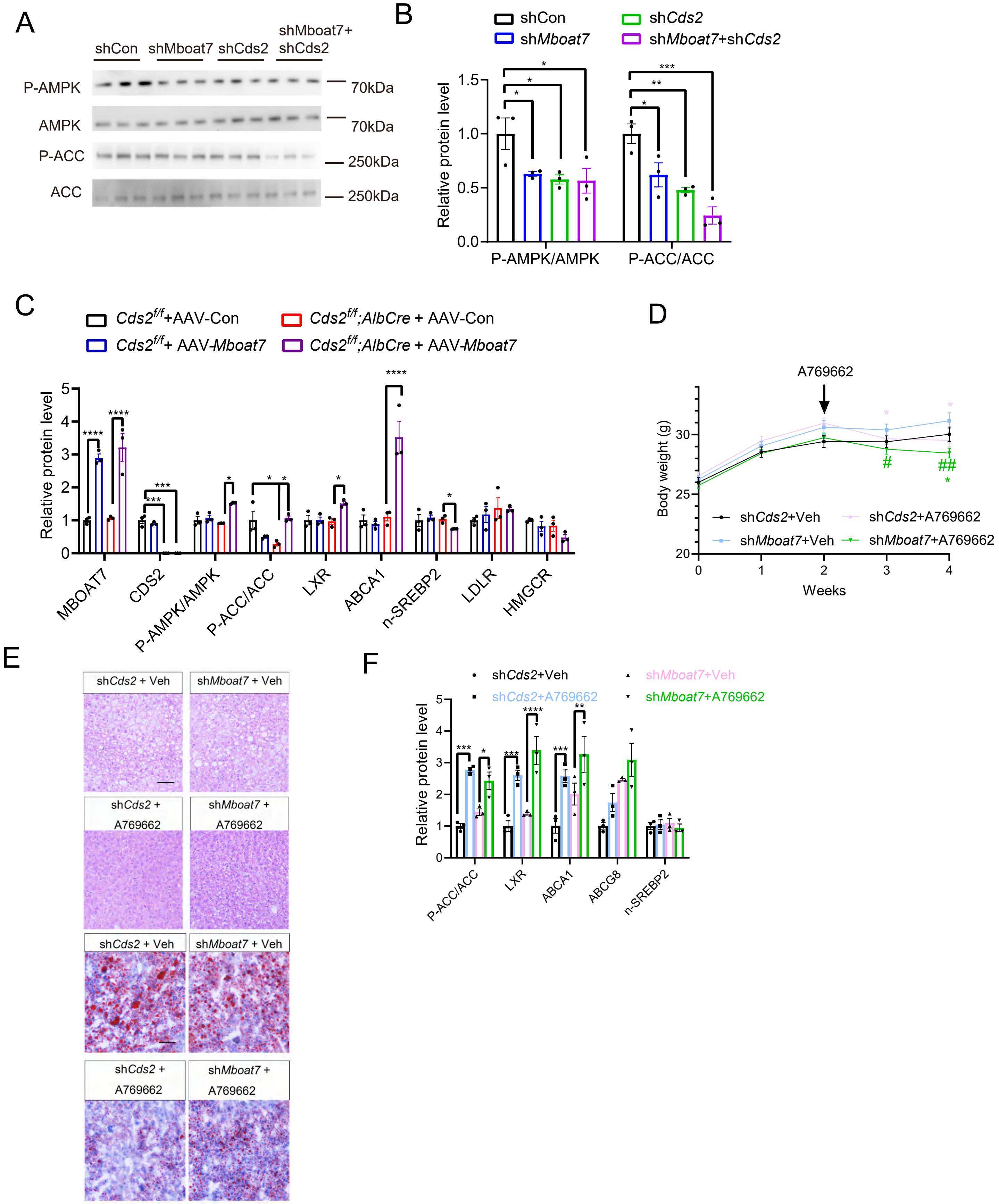

A
shCon
shMboat7
shCds2
shMboat7+
shCds2
P-AMPK
AMPK
P-ACC
ACC
70kDa
70kDa
250kDa
250kDa
B
shCon
shMboat7
shCds2
shMboat7+shCds2
Relative protein level
P-AMPK/AMPK
P-ACC/ACC
C
Cds2f/f+AAV-Con
Cds2f/f;AlbCre + AAV-Con
Cds2f/f+ AAV-Mboat7
Cds2f/f;AlbCre + AAV-Mboat7
Relative protein level
MBOAT7
CDS2
P-AMPK/AMPK
P-ACC/ACC
LXR
ABCA1
n-SREBP2
LDLR
HMGCR
D
A769662
Body weight (g)
Weeks
shCds2+Veh
shCds2+A769662
shMboat7+Veh
shMboat7+A769662
E
shCds2 + Veh
shMboat7 + Veh
shCds2 + A769662
shMboat7 + A769662
shCds2 + Veh
shMboat7 + Veh
shCds2 + A769662
shMboat7 + A769662
F
shCds2+Veh
shMboat7+Veh
shCds2+A769662
shMboat7+A769662
Relative protein level
P-ACC/ACC
LXR
ABCA1
ABCG8
n-SREBP2
